# Characterization of a heterogenous activated B cell compartment arising early after antigen exposure preceding long-lived memory B cell formation

**DOI:** 10.1101/2024.12.22.629610

**Authors:** L Fernandez Blanco, LH Kuijper, LYL Kummer, NJM Verstegen, A Bos, M Claireaux, MC Duurland, T Jorritsma, M Steenhuis, G Kerster, JJ Garcia Vallejo, MJ van Gils, PJ van Dam, EW Stalmam, L Wieske, L Boekel, GJ Wolbink, SW Tas, T Rispens, TW Kuijpers, F Eftimov, A ten Brinke, SM van Ham, T2B! Immunity against SARS-CoV-2 study group

## Abstract

Once formed, plasma cells and memory B cells (MBCs) are difficult to eradicate, posing a problem in the context of unwanted antibody responses. Characterizing early B cell differentiation stages after antigen encounter is thus crucial to target and prevent unwanted antibody formation. Here, we unravelled in-depth antigen-specific B cell responses longitudinally after SARS-CoV-2 mRNA vaccination in healthy individuals using multiparameter spectral flow cytometry. The early antigen-specific B cell response was dominated by spike-specific IgG^+^ CD27^+^ CD71^+^ activated B cells (ActBCs), previously assigned as germinal center-derived and DN2 extrafollicular B cells. Within the early IgG^+^ ActBC compartment, six distinct clusters were identified with specific contraction dynamics, whereby some of these clusters were more closely related to pre-ASCs and others more to long-lived MBCs. Some of the highly contracting ActBC clusters expressed CD11c, a marker previously used to define atypical B cells. The transient presence of different ActBC clusters could also be observed in total B cells when gated in an antigen-independent manner. Our results thus delineate the early stages of the antigen-specific B cell response, with a further dissection of the CD71^+^ ActBC compartment. Detection of ActBC clusters early after antigen encounter in total B cells opens avenues for future evaluation of their potential to serve as a proxy for antigen-reactive B cells in autoimmunity or other unwanted B cell responses.

## Introduction

Vaccination is an important strategy to battle infectious diseases by inducing long-lasting cellular and humoral immune responses. Antibodies protect against new and recurring pathogens and are generated by antibody-secreting cells (ASCs) induced by *de novo* naïve B cell differentiation or upon reactivation of memory B cells (MBCs). Both T cell-independent or T cell-dependent B cell differentiation routes have been identified (1–3). Unfortunately, antibody responses can also be detrimental in the context of autoimmunity, alloimmunization or reactivity against therapeutic biologics. Once established, long-lived ASCs and MBCs are difficult to eradicate. Therefore, identification of ASC and MBC precursors is warranted and knowledge on their differentiation status is much desired to design strategies to optimize vaccination responses or, conversely prevent unwanted antibody formation.

Upon primary vaccination, a first rapid antibody response is induced. The extrafollicular (EF) pathway is a fast response that leads to formation of short-lived plasmablasts (PBs) and early MBCs (3, 4). Besides this EF response, germinal centers (GC) are formed in secondary lymphoid organs where antigen-stimulated B cells proliferate and undergo class-switch recombination and somatic hypermutation (SHM) in the dark zone and repeatedly cycle to the light zone, reencounter antigen and receive renewed T cell help (4, 5). This eventually allows for selection of B cells with improved BCR affinity for the antigen and leads to generation of long-lived MBCs and ASCs. From the latter, long-lived plasma cells (PCs) are established upon migration to the bone marrow (6–8). Using high-output techniques, recent studies have identified diverse subphenotypes of circulatory MBCs based on different surface markers (9–11). However, to what extent these different phenotypes share developmental trajectories or reflect different subfunctions, especially in the context of vaccination responses, requires further in-depth research. In addition, the exact decision point and B cell stage in which bifurcation between MBC or ASC phenotype occurs remains to be elucidated. Recent studies into this subject have investigated activated B cells (ActBCs), a population of antigen-specific B cells leaving the GC, which are phenotypically and functionally different from ASCs (10, 12–14). Ellebedy and colleagues identified CD71 as a marker for recently activated B cells (12). Lau and colleagues defined low expression of CD21 as a marker of CD27^+^ B cells that recently emigrated from the GC and are potential PC precursors (13). So-called ActBCs peak in circulation shortly after immunization or infection and drop again over time, while at the same time circulating MBCs start to expand (11–15). Based on their phenotypic definition, we recently reported that ActBCs may exhibit diverse fates with some feeding the resting MBC compartment and others sustaining the long-lived ASCs (11, 16). Both classical ASC and MBC subsets have been widely studied in context of vaccination and infection (12, 15, 17, 18). However, a deeper understanding and characterization of the ActBC compartment is lacking and much needed to monitor early stages of B cell differentiation after antigen exposure and to interfere with it before ASC and MBC formation has occurred in cases of unwanted B cell responses.

In recent years, the EF B cell response has acquired more attention, with a specific focus on Tbet^+^ CD11c^+^ CD21^-/lo^ cells, commonly referred to as atypical B cells, in the context of autoimmunity, chronic infections or vaccination (14, 19, 20). The heterogeneous nature of this population and its contribution to both normal and aberrant immune responses have led to inconsistencies in its classification and nomenclature. CD11c^+^ B cells are mainly categorized by presence or absence of CD27 expression: IgD^+^ CD27^-^ ‘activated-naïve’ B cells and IgD^-^ CD27^-^ ‘double-negative 2’ B cells (DN2), identified as part of the EF response, and CD27^+^ ‘age-associated B cells’ (ABCs), which formation also remains unclear as they could have an EF origin or be GC-derived an acquire CD11c upon reactivation (21). Both populations, DN2 and ABCs, are thought to be poised for ASC differentiation, although their specific role and contribution within the B cell differentiation process remains unclear (19, 22–25). Although Tbet^+^ CD11c^+^ CD21^-/lo^ B cells comprise only a small part of the total CD19^+^ B cell compartment, this population expands vastly after vaccination in humans (13, 17, 24, 26). We recently reported DN2 and ABCs to be among the early B cell responding populations after antigen exposure and even to be main contributors to the S (Spike)-specific response seven days after Severe Acute Respiratory Syndrome Coronavirus 2 (SARS-CoV-2) vaccination while contracting over time (24). It remains to be established how CD27^+^ CD11c^+^ CD21^-/lo^ B cells and IgG^+^ ActBCs relate to each other and to MBCs versus PC formation.

In this study, we performed in-depth characterization of antigen-specific B cells to investigate the dynamics and early stages of human B cell differentiation in COVID-19 naïve individuals after SARS-CoV-2 mRNA- 1273 vaccination. We studied the kinetics of S- and receptor binding domain (RBD)-specific B cell populations and compared this to antigen-specific Influenza hemagglutinin (Influenza-HA), fusion glycoprotein from Respiratory Syncytial Virus (RSV-F) and Tetanus Toxoid (TT) B cell populations which were encountered longer ago. ActBCs and EF B cells could be identified early after vaccination followed by establishment of resting MBCs six months after antigen exposure. Overall, several distinct ActBC phenotypes were identified, characterized by two clear dynamics: a pronounced and significant contraction between day seven and six months post-vaccination in contrast to clusters showing no significant contraction and more inter-donor variation in contraction dynamics. Remarkably, a major proportion of the highly dynamic ActBC clusters expressed CD11c, which underlines that the distinction between EF and GC-derived B cells remains to be redefined. The dynamic IgG^+^ CD11c^+^ ActBC clusters were identified in total CD19^+^ B cells without the use of antigen probes shortly after vaccination.

## Methods

### Study design and participants

This study is part of a prospective observational cohort study in the Netherlands, called the Target-to-B! SARS-CoV-2 vaccination study including healthy individuals and patients with autoimmune disease. Full study details have been previously published (27). The study was approved by the medical ethical committee (NL74974.018.20 and EudraCT 2021-001102-30, local METC number: 2020_194) and registered at the Dutch Trial register (trial ID: NL8900). All participants provided written informed consent. Participants were vaccinated between April 2021 and October 2021 with the mRNA-1273 (Moderna) vaccine. The primary vaccination was given at a six-weeks interval, conform the Dutch national vaccination campaign guidelines at the time. Peripheral blood was collected by venipuncture seven days and six months post second vaccination. Peripheral blood mononuclear cells (PBMCs) were isolated within 12 hours and frozen at -80 ⁰C. In total 104 participants of the Target-to-B study cohort were used for the analysis of antigen-specific B cells, including patients with rheumatoid arthritis (RA) or inflammatory bowel disease (IBD) treated with immunomodulating therapies, as well as RA and IBD disease controls and healthy controls. Adult healthy participants were actively recruited at the Reade center of Rheumatology in Amsterdam, the Netherlands, and were friends or family members of patients willing to participate as healthy controls. In the current study we focused the analysis only on the healthy controls. Exclusion criteria were active or previous autoimmune-, oncological- or hematological disease; current or previous treatment with systemic medication in the past year, pregnancy and previous SARS-CoV-2 infection (as evidenced by positive anti-RBD antibodies before first vaccination and/or self-reported positive PCR and/or positive anti-nucleocapsid protein (NCP) antibodies throughout the study period).

### Probe design

All recombinant, soluble proteins, including SARS-CoV-2 S-2P, RBD, Influenza A hemagglutinin (H1N1pdm2009), RSV prefusion stabilized F (DS-Cav1) were produced as described before (11). After purification, proteins were biotinylated with a BirA500 551 biotin-ligase reaction kit (Avidity). TT was purchased from 552 Creative Biolabs (Vcar-Lsx003). NCP was produced as described before (28). TT and NCP were unspecifically biotinylated using EZ-Link Sulfo-NHS-LC553 Biotinylation Kit (Thermo Fisher).

### Spectral flow cytometry of antigen-specific B cells

PBMCs were thawed in IMDM (Lonza) with 10% FCS (Bodinco BV). 10 x 10^6^ PBMCs were enriched for B cells using CD3 T cell depletion via the EasySep™ Human CD3 Positive Selection Kit II (StemCell Technologies) according to the manufacturer’s instruction. To stain antigen-specific B cells, biotinylated protein antigens SARS-CoV-2 S, RBD, NCP, Influenza-HA, RSV-F and TT were individually multimerized with fluorochrome-conjugated streptavidin in a 2:1 molar ratio at 4°C for 1 hour. Next, 10% D-biotin (GeneCopoeia) was added and incubated at 4°C for at least 30 minutes to block free sites on the streptavidin and reduce cross-reactivity among biotinylated protein antigens. All biotinylated antigens were conjugated to two fluorochromes, except RBD, which was conjugated with a single fluorochrome and gated as part of S1 (**Table S1**). A 31-color spectral cytometry panel, including the 6 antigen proteins, was designed to phenotype antigen-specific B cells (**Table S2**). Enriched B cell samples were first stained using Live/Dead Fixable Blue Stain Kit (Invitrogen) in PBS at 4°C for 30 min. Subsequently, cells were washed twice with staining buffer (PBS supplemented with 1% BSA and 1mM EDTA) and stained with anti-IgG for 10 min at 4°C and immediately after with the rest antibody mix and antigen multimer cocktail for 30 min at 4°C. Next, cells were washed twice with PBS, fixed with 1% paraformaldehyde for 10 minutes at 4°C and washed twice again with staining buffer. Data were acquired on Cytek Aurora 5L spectral cytometer using SpectroFlo® software (Cytek Biosciences).

### Spectral flow cytometry data pre-processing

After spectral unmixing (SpectroFlo v3.0.1, Cytek), FCS files were loaded into OMIQ analysis software (www.omiq.ai). Initial gating was performed to select for single, live and CD19^+^ DUMP^-^ (CD3, CD4,

CD16, CD56) cells. PeacoQC was run to detect and remove flow cytometry anomalies in both signal acquisition and dynamic range (29). To exclude batch effects, all data were normalized using Cytonorm for 8 markers (CD19, CD20, CD24, CD38, CD45RB, HLA-DR, IgD, IgM) based on a reference sample which was measured in each batch (30). Subsequently, negative gates were set for each combination of probes, after which antigen-specific B cells were further positively gated based on their specific combinations of fluorochrome-conjugated streptavidin (**Figure S1**). For downstream analysis, data were subsampled to include only all detected antigen-specific B cells (n = 450.000)

### Dimensionality reduction and FlowSOM clustering

For UMAP visualization and FlowSOM clustering (xdim 16, ydim 16), input consisted of a selection of 12 markers (CD20, CD21, CD27, CD24, CD38, CD138, CD11c, CD45RB, IgA, IgD, IgM, IgG). Antigen-specific B cell data from 104 participants of the T2B study cohort were used. FlowSOM clusters were manually merged and annotated based on UMAP visualization, heatmap analysis of median marker expression between clusters and manual gating of specific marker combinations (**Table S3**).

### ELISA assays

Serum samples were collected at different time points using venipuncture and home-based finger prick kits to detect the presence of RBD and NCP-specific antibodies. An in-house developed anti-RBD IgG ELISA was used as described before (28, 31). First, anti-RBD IgG levels were measured with a quantitative ELISA and expressed as arbitrary units per milliliter (AU/mL). Second, a semi-quantitative total antibody bridging ELISA to detect NCP antibodies was used to identify SARS-CoV-2 infection after first vaccination (31).

### Statistical analysis

Statistical analysis, graphs, heatmaps and stacked bar graphs were made using Rstudio (version 4.1.1) and GraphPad Prism 9.1.1. Computational analysis of data to generate heatmaps was performed using the Spectre R package (32). Statistical significance was assessed using Wilcoxon signed-rank test for paired data and p-values were corrected for multiple comparison using *post-hoc* Bonferroni-Holm’s test. P-values of <0.05 were considered significant.

## Results

### Dynamics of the antigen-specific B cell compartment following mRNA vaccination

To assess the dynamics of antigen-specific B cell differentiation after recent *de novo* antigen exposure, a cohort including 18 SARS-CoV-2 mRNA vaccinated healthy individuals was selected, consisting of 10 females and 8 males with a median age of 48 years old (SD=10) (**Table 1**). All individuals received two mRNA-1273 (Moderna) vaccinations and were confirmed SARS-CoV-2 naïve before vaccination. PBMCs were collected prior to first vaccination (V1pre, n=18), seven (V2D7, n=18) and 187 days (V2M6, n=14) after second vaccination (**Figure 1A**). All individuals seroconverted after primary vaccination and remained seropositive up to 6 months post vaccination (**Figure S1A, Table 1**). High-dimensional analysis was performed using a 31-color spectral flow cytometry panel (**Table S1, 2**), including five fluorophores for combinatorial antigen labelling (S, RBD, NCP, HA, RSV-F, TT), 21 B cell surface markers and the four major immunoglobulin isotypes. NCP labelling was used to confirm absence of previous SARS-CoV-2 infection at baseline and during the study. Of note, during the period of sampling, influenza and RSV infections were uncommon due to lockdown and TT (DTP) revaccination was unlikely due to travel restrictions. S- and RBD-specific B cells made up on average 0.4% and 0.1% of total CD19^+^ B cells, respectively, at day seven and remained constant up to six months following vaccination, albeit with a declining trend for RBD (**Figure 1B, Figure S1B-C**). Over time, an increase in S/RBD ratio was observed (**Figure S1D**), indicating a shift of the SARS-CoV-2 specific B cell receptor repertoire over time towards non-RBD epitopes, in line with previously reports by us and others (33, 34).

**Figure 1.**
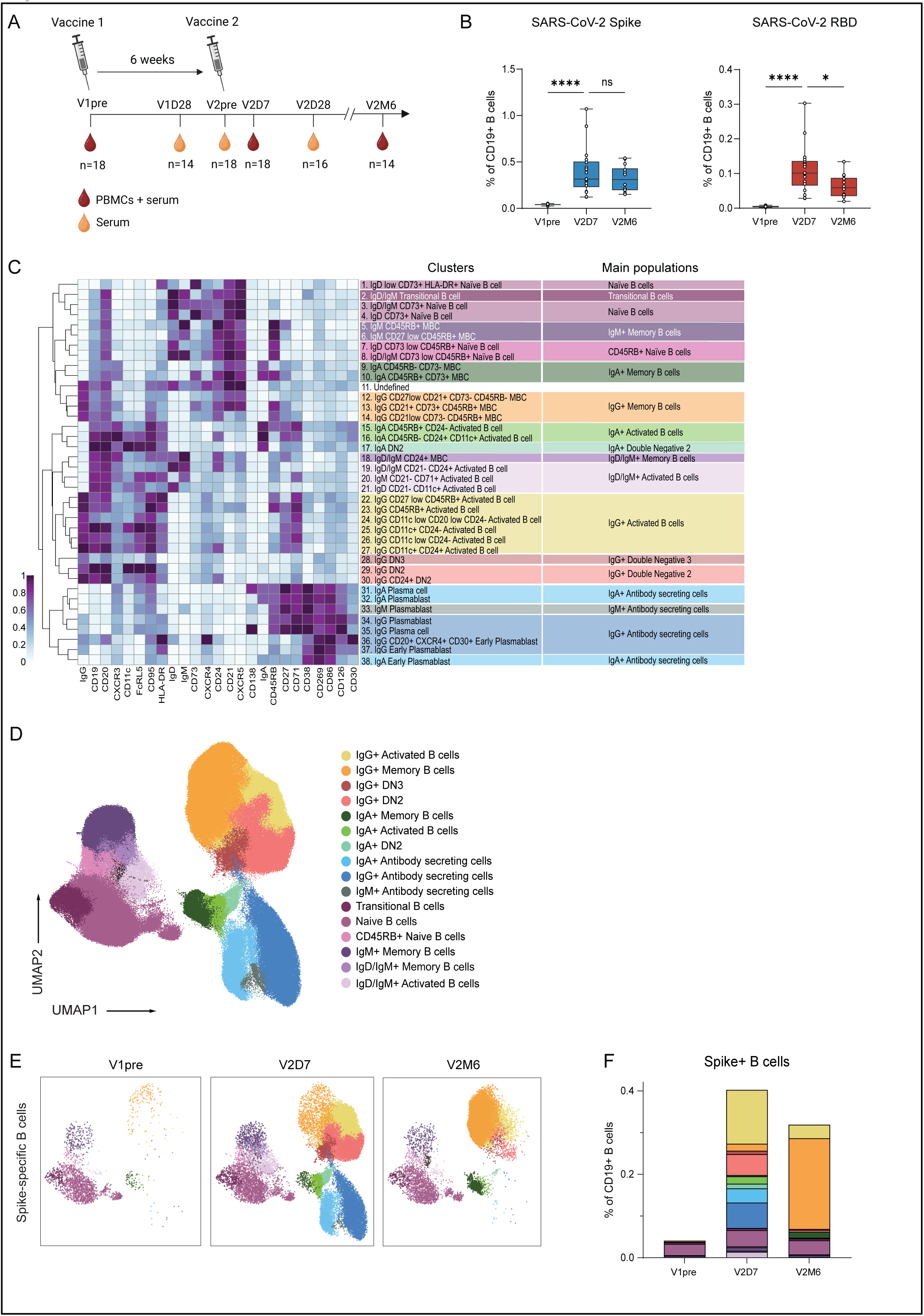
Longitudinal dynamics of the antigen-specific B cell compartment following SARS-CoV-2 mRNA vaccination. **A.** Study design and timeline with SARS-CoV-2 mRNA vaccination administration and peripheral blood and serum collection. **B.** Percentage of S- and RBD-specific B cells of total CD19^+^ B cells at V1pre (n=18), V2D7 (n=18) and V2M6 (n=14). **C.** Heatmap of the 38 B cell clusters and 16 B cell main populations after FlowSOM clustering. **D.** UMAP projection of all antigen-specific B cells colored by the 16 main populations. **E.** UMAP projection of S-specific B cells per time point. **F.** Stacked bar graph representing the mean proportion of S^+^ B cells (% of CD19^+^) for the 16 main populations per time point. Statistical significance was assessed using Wilcoxon signed-rank test for paired data and p-values were corrected for multiple comparison using *post hoc* Bonferroni-Holm’s test. (\**P* < 0.05, \*\**P* < 0.01, \*\*\**P* < 0.001, \*\*\*\**P* < 0.0001).

**Table 1.**

|  | Healthy controls (n=18) |
| --- | --- |
| **Age in years, mean (SD)** | 48 (10) |
| **female sex, n (%)** | 10 (55%) |
| **vaccine type, n (%)** |  |
| mRNA-1273 (Moderna) | 18 (100%) |
| **Time between V2pre and V2post 7d, in days, median (IQR)** | 7 (0) |
| **Time between V2pre and V2post 6m, in days, median (IQR)** | 187 (7) |
| **Seroconversion after second vaccination\*, n (%)** | 18 (100%) |
| Anti-RBD IgG titer, median (IQR) | 234 (110) |

To capture the phenotypic complexity of the antigen-specific B cell response, uniform manifold approximation projection (UMAP) dimensionality reduction was used to visualize all SARS-CoV-2 (S, RBD) and other (TT, RSV-F, HA) antigen-specific CD19^+^ B cells, and subsequently FlowSOM clustering was performed on a selection of core markers (CD20, CD21, CD27, CD138, CD38, CD24, CD45RB, CD11c, IgM, IgA, IgG, IgD) on all antigen-specific B cells. Clustering analysis revealed 38 distinct clusters that could be manually grouped into 16 overarching B cell populations, or metaclusters, based on expression of 25 surface markers (**Figure 1C-D, Figure S1E, Table S3**). Among the 16 main populations, transitional B cells, naïve B cells, MBCs, ActBCs and ASCs were identified. The naïve compartment (IgD^+^/IgM^+^ CD27^-^ CD21^+^), as visualized in the UMAP, contained two different populations based on CD45RB and CD73 expression. Of note, the naïve compartment is not truly naïve and could be better defined as antigen-reactive. Adjacent to the naïve B cells in the UMAP, IgD/IgM^+^ MBCs and ActBCs (CD27^-^ CD21^-^) were identified. The upper area of the UMAP depicts the IgG compartment, including IgG^+^ MBCs, ActBCs (CD27^+/lo^ CD38^lo^ CD21^lo^ CD71^+^ CD95^hi^), IgG^+^ DN2 (IgD^-^ CD27^-^ CD21^-^ CD11c^+^), and IgG^+^ DN3 (IgD^-^

CD21^-^ CD27^-^ CD11c^-^) B cell populations. Additionally, MBCs, ActBCs and DN2 populations were also observed for the IgA isotype. Lastly, the ASC compartment, annotated based on low CD19 and CD20 and high CD38 expression, represented all three BCR isotypes. Seven days after second vaccination, the majority of the S- and RBD-specific B cell response comprised of IgG^+^ ActBCs, followed by IgG^+^ and IgA^+^ ASCs and IgG^+^ DN2. After six months, IgG^+^ MBCs dominated the S- and RBD-specific B cell response while ActBCs, DN2 and ASCs had contracted (**Figure 1E-F, Figure S1F-G**). Given the fragility of ASCs and since frozen cells were used, it is likely that we underestimated the frequency of the ASC compartment, although we were able to identify eight different clusters (17, 35). The IgG^+^ ASCs consisted of PCs (CD27^hi^ CD38^hi^ CD138^+^), PBs (CD27^hi^ CD38^hi^) and early PBs (CD27^lo^ CD38^+^) clusters, with IgG^+^ PB as the most abundant cluster. All clusters demonstrated similar dynamics of expansion early after vaccination and contraction six months after (**Figure 2**). Interestingly, a minor IgG^+^ CD20^+/int^ CXCR4^+^ CD30^+^ cluster 36 was identified. Similar to pre-plasmablast clusters, cluster 36 was CD38^+^ CD27^lo^ and had high CD269 expression, but was unique in high CD30 and intermediate CD20 expression (**Figure 2A-B**).

**Figure 2.**
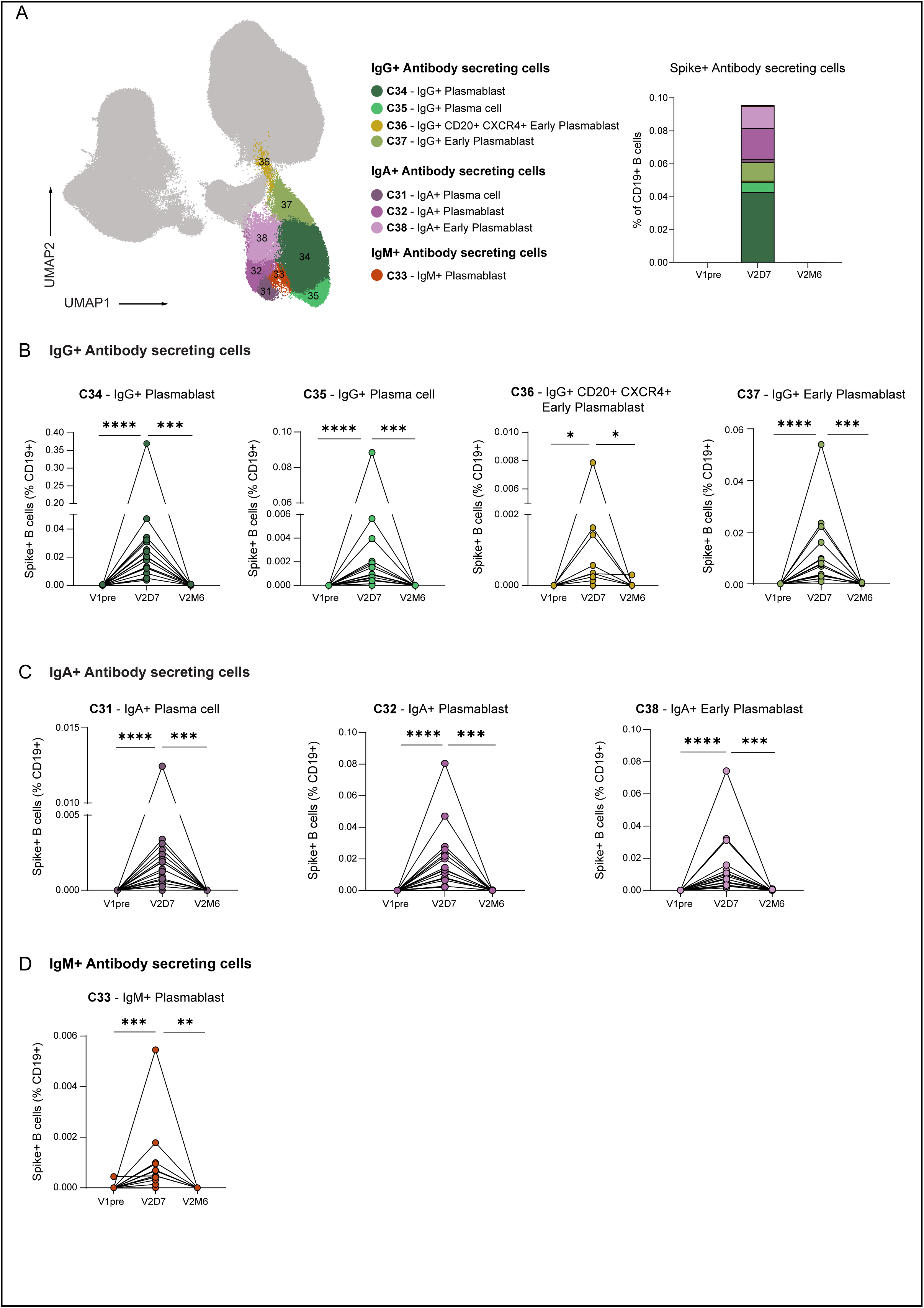
Dynamics of the ASC compartment after SARS-CoV-2 mRNA vaccination. **A.** UMAP projection of IgG^+^, IgA^+^ and IgM^+^ ASC clusters. Stacked bar graph representing the mean proportion of S^+^ ASCs (% of CD19^+^) per cluster over time. **B.** Dynamics of IgG^+^ ASC clusters. **C.** Dynamics of IgA^+^ ASC clusters. **D.** Dynamics of IgM^+^ ASC cluster. V1pre (n=18), V2D7 (n=18) and V2M6 (n=14). Statistical significance was assessed using Wilcoxon signed-rank test for paired data and p-values were corrected for multiple comparison using *post hoc* Bonferroni-Holm’s test. (\**P* < 0.05, \*\**P* < 0.01, \*\*\**P* < 0.001, \*\*\*\**P* < 0.0001).

Together, these data show that the ActBC, DN2 and ASC compartments dominated the antigen-specific response seven days after second vaccination and contract six months after, while the MBC compartment is dominant six months after vaccination.

### Classical memory B cells can be subphenotyped beyond CD27 expression using CD21, CD73 and CD45RB

To further explore the dynamics and phenotypic relationships of S-specific IgG^+^ B cells after vaccination, the IgG^+^ B cell compartment was further investigated **(Figure 3A**). This compartment consists of classical MBCs, ActBCs and DN B cells. Classical IgG^+^ MBCs are generally defined as CD27^+^ CD38^-^ B cells. However, in our clustering, the absence of CD38 and CD71, reflecting a non-activated phenotype, were the main discriminative markers as CD27 showed an intermediate expression. Among classical MBCs, three different clusters (cluster 12, 13 and 14) could be distinguished based on unique dynamic frequencies over time. Cluster 12 and 13 dominated the S-specific response six months after second vaccination, while cluster 14 expanded less between day seven and six months after vaccination (**Figure 3B-C**). We further analysed these different memory phenotypes in the HA-, TT- and RSV-F-specific B cell compartments, which have been established months to years ago. Cluster 13 dominated the HA-, TT- and RSV-F-specific B cell compartment, while cluster 12 and 14 were only marginally present, suggesting that cluster 13 is the most long-lived resting MBC cluster **(Figure 3B-C**). Cluster 13 expresses CD21, CD45RB and also a vast majority of the cells expresses CD73, in line with previous observations by us and others (11, 35). CD73 is a marker for metabolic quiescence in resting MBCs, which further substantiates cluster 13 long-lived characteristic. Regarding CD45RB, in our data, it does not show a significant discriminative power between two branches of MBCs as it has been recently described (10). Although cluster 12 showed a similar resting dynamic as cluster 13 and both clusters were CD21^+^, cluster 12 expressed low levels of CD27, CD73 and CD45RB (**Figure S2**). Moreover, since cluster 12 was not dominant in the TT-, HA- and RSV-F-specific compartment, this cluster appears less long-lived compared to cluster 13. Alternatively, cluster 12 could be a specific MBC phenotype for vaccination responses. Lastly, cluster 14 was CD45RB^+^ but CD73^-^ and displayed a more activated phenotype, concomitant with low CD21 expression. This phenotype in combination with less profound expansion between days seven and six months after vaccination compared to cluster 12 and 13, suggests that cluster 14 may be a more intermediate population between ActBCs and MBCs (**Figure 3C**).

**Figure 3.**
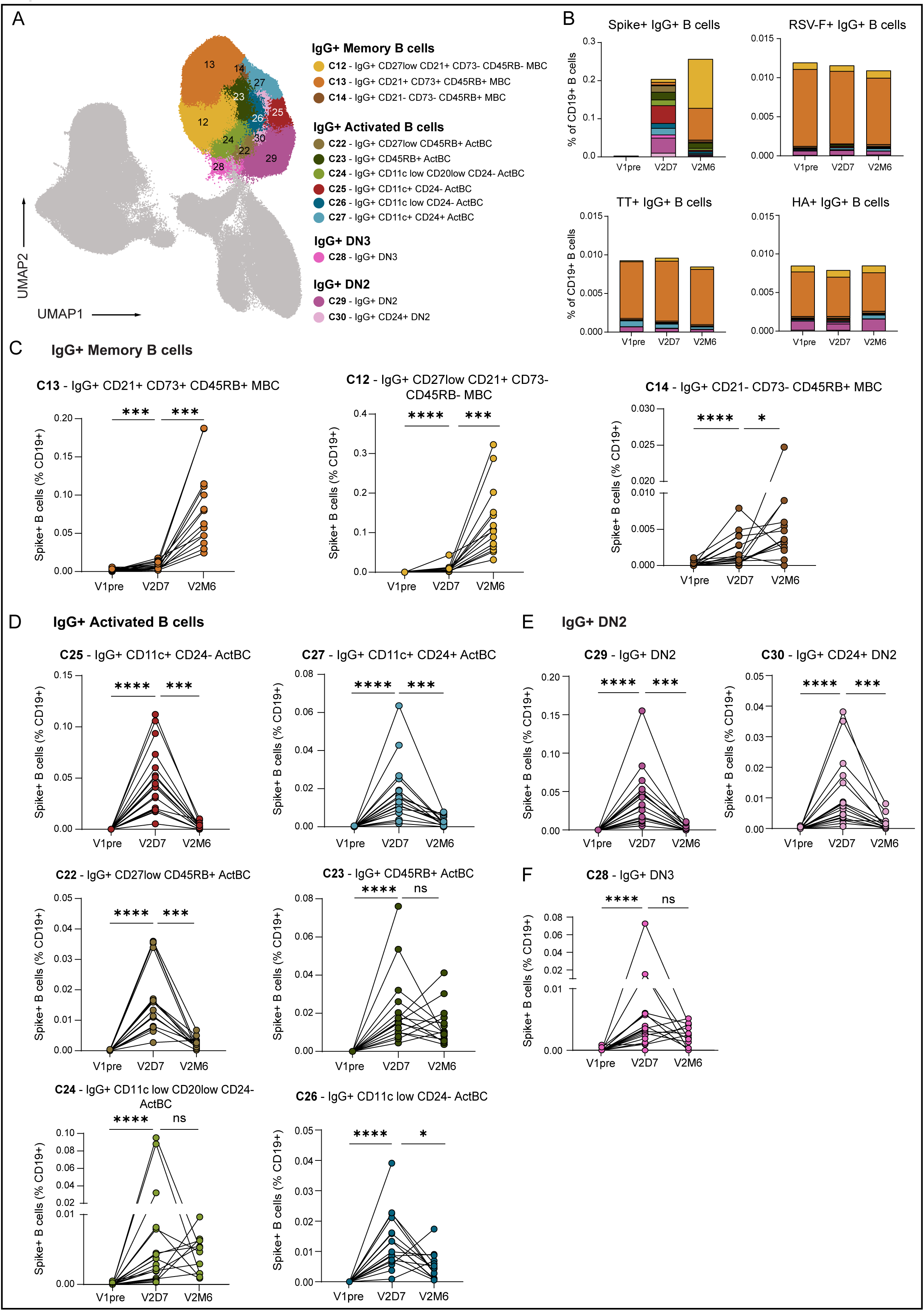
Dynamics of IgG+ B cell compartment after SARS-CoV-2 mRNA vaccination. **A**. UMAP projection of IgG^+^ B cells including MBCs, ActBCs and DN colored per cluster. **B.** Stacked bar graph representing the mean proportion of S^+^, TT^+^, Influenza-HA^+^ and RSV-F^+^ IgG^+^ B cells as percentage of CD19^+^ B cells over time. **C.** Dynamics of IgG^+^ MBC clusters as percentage of S^+^ B cells (% of CD19^+^). **D.** Dynamics of IgG^+^ ActBC clusters as percentage of S^+^ B cells (% of CD19^+^). **E.** Dynamics of IgG^+^ DN2 clusters as percentage of S^+^ B cells (% of CD19^+^). **F**. Dynamics of IgG^+^ DN3 cluster as percentage of S^+^ B cells (% of CD19^+^). V1pre (n=18), V2D7 (n=18) and V2M6 (n=14). Statistical significance was assessed using Wilcoxon signed-rank test for paired data and p-values were corrected for multiple comparison using *post hoc* Bonferroni-Holm’s test. (\**P* < 0.05, \*\**P* < 0.01, \*\*\**P* < 0.001, \*\*\*\**P* < 0.0001).

To conclude, expression of CD73, CD21 and CD45RB defines the most long-lived resting MBC compartment (cluster 13). Cluster 12 and 14 are dynamically and phenotypically related to these cells, but exhibit reduced longevity as they are much less present long after antigen exposure.

### The IgG Activated B cell compartment is a heterogeneous population with diverse dynamics similar to both DN B cells and the MBC compartment

Considering the enrichment of the S-specific B compartment with IgG^+^ ActBCs and DN B cells early after vaccination (11, 14, 37), we looked deeper into the phenotype and dynamics of these compartments following vaccination (**Figure 3A**). The overall IgG^+^ ActBC compartment was defined based on CD27 and high CD71 expression and low expression of CD21 and CD38, whereas DN B cells were characterized by lack of CD27, CD21 and IgD. FlowSOM clustering identified six different IgG^+^ ActBC clusters (cluster 22 and 27), based on differences in CD45RB, CD24 and CD11c expression, reflecting a high level of heterogeneity in this compartment (**Figure 3A, Figure S2**). Additionally, clustering revealed three different DN clusters distinguished by CD11c and CD24 expression, two defined as IgG^+^ DN2 (cluster 29 and 30) and one IgG^+^ DN3 (cluster 28) population. ActBC clusters 25, 27 and 22, although prominently present seven days after second vaccination, contracted almost completely six months after, indicative of an early, activated phenotype that is not maintained over time (**Figure 3D**). ActBC cluster 26 also contracted over time, but less significantly. Remarkably, cluster 25 and 27 expressed CD11c, defining an antigen-specific ActBC population co-expressing CD71 and CD11c (**Figure S2**). IgG^+^ DN2, which also express CD11c but lack CD27 and CD21 expression, displayed similar dynamics over time following vaccination (**Figure 3E**). Interestingly, ActBC cluster 22 expressed low levels of CD27 and was located closely to the DN B cells in the UMAP. On the other hand, ActBC clusters 23 and 24 did not contract significantly over time, with even expansion of these clusters in some individuals, reflecting strong inter-donor variation, between day seven and month six after vaccination (**Figure 3D**). Fitting with the less contracting profile over time, cluster 23 was located closely to the long-lived MBC compartment in the UMAP (**Figure 3A**). In addition, cluster 23 moderately expressed CD21, indicative of an intermediate phenotype between ActBCs and MBCs, as also previously suggested for cluster 14 (**Figure 3D).** Interestingly, CD24, a marker recently described by our group to be expressed on activated MBCs in their trajectory to become resting MBCs, was expressed in the more dynamic cluster 27 (11). IgG^+^ DN3 B cells, which resemble DN2 in their CD27^-^ CD21^-^ phenotype but lack CD11c expression, were less dynamic over time and were located near both, dynamically active ActBCs but also near the more stable ActBC cluster 24 and the pre-PB cluster 36 in the UMAP (**Figure 3F**). Interestingly, both cluster 24 and the DN3 showed less CD20 expression than other B cell populations. Collectively, these findings suggest that IgG^+^ ActBCs and DN B cells show a partial phenotypic overlap and are established early after antigen exposure, while quickly contracting in peripheral blood over time.

### Activated B cell heterogeneity is also evident for B cells expressing IgM or IgA

To further understand the activation profile of the antigen-specific B cell compartment after vaccination, we investigated whether the activation markers identified for the IgG^+^ B cell compartment were also indicative of IgA^+^ or IgM^+^ B cell activation. Overall, as expected, IgA^+^ B cells were less abundant than IgG^+^ B cells in the S-specific response following vaccination, but were similar in phenotype and dynamic behavior. The IgA^+^ B cell compartment included MBCs, ActBCs and DN2 (**Figure 4A)**. FlowSOM clustering identified two IgA^+^ MBC clusters (cluster 9 and 10), which were mainly present later in the vaccination response. Similar to IgG^+^ MBC cluster 13, IgA^+^ MBC cluster 10 exhibited a CD21^+^ CD73^+^ CD45RB^+^ phenotype, corresponding with the long-lasting MBC compartment (**Figure 4B**). Furthermore, two IgA^+^ ActBCs clusters (cluster 15 and 16) were identified. Both clusters peaked at day seven and contracted six months later, but differed phenotypically in CD45RB, CD24 and CD11c expression. IgA^+^ cluster 15 was CD45RB^+^ CD24^-^ CD11c^-^, which phenotypically resembled cluster 23 in the IgG compartment. However, this cluster was short-lived as opposed to cluster 23. IgA ^+^ cluster 16 was with CD45RB^-^ CD24^+^ CD11c^+^ expression phenotypically similar to IgG ^+^ cluster 27 (**Figure 4C-D**), indicating that both IgG ^+^ and IgA^+^ ActBCs can concomitantly express CD71 and CD11c.

**Figure 4.**
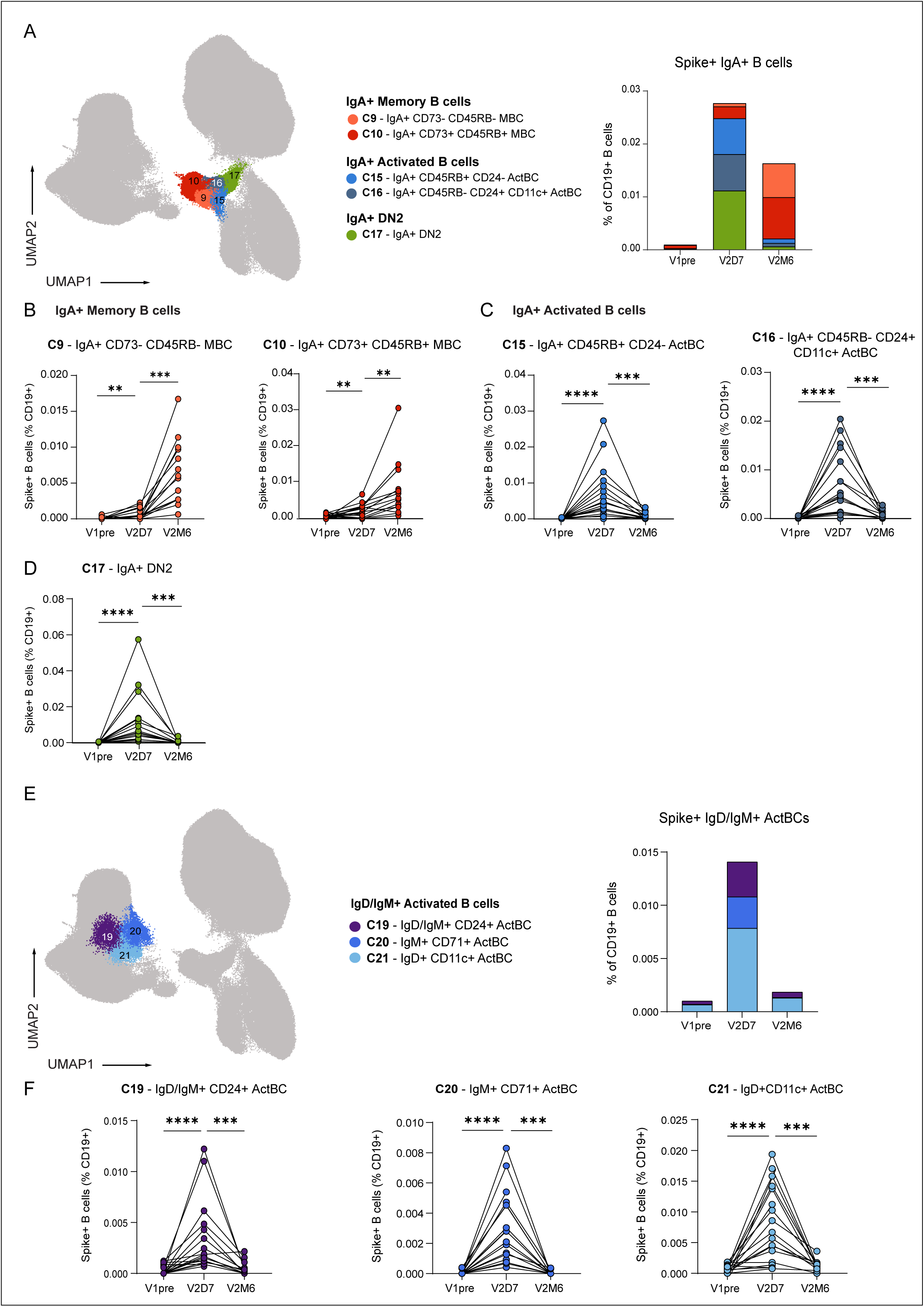
Dynamics of IgA+ and IgD/IgM+ B cell compartment after SARS-CoV-2 mRNA vaccination. **A.** UMAP projection of IgA^+^ B cells including MBCs, ActBCs and DN2 colored per cluster. Stacked bar graph representing the mean proportion of S^+^ IgA^+^ B cells (% of CD19^+^) per cluster over time. **B.** Dynamics of IgA^+^ MBC clusters as percentage of S^+^ B cells (% of CD19^+^). **C.** Dynamics of IgA^+^ ActBC clusters as percentage of S^+^ B cells (% of CD19^+^). **D.** Dynamics of IgA^+^ DN2 clusters as percentage of S^+^ B cells (% of CD19^+^). **E.** UMAP projection of IgD/IgM^+^ ActBCs including MBCs, ActBCs and DN2 colored per cluster. Stacked bar graph representing the mean proportion of S^+^ IgD/IgM^+^ ActBCs (% of CD19^+^) per cluster over time. **F.** Dynamics of IgD/IgM^+^ ActBCs clusters as percentage of S^+^ B cells (% of CD19^+^). V1pre (n=18), V2D7 (n=18) and V2M6 (n=14). Statistical significance was assessed using Wilcoxon signed-rank test for paired data and p-values were corrected for multiple comparison using *post hoc* Bonferroni-Holm’s test. (\**P* < 0.05, \*\**P* < 0.01, \*\*\**P* < 0.001, \*\*\*\**P* < 0.0001).

In addition to IgA, we identified three IgD^+^ and/or IgM^+^ CD21^-^ CD27^-^ B cell clusters, located next to the naïve compartment but displaying similar dynamics to IgG^+^ and IgA^+^ ActBCs and DN2 B cells: high induction at day seven and contraction at six months. The three clusters additionally expressed CD24 (cluster 19), CD71 (cluster 20) or CD11c (cluster 21) (**Figure 4E-F**). Cluster 21 (IgD^+^ CD11c^+^ ActBCs), previously referred to as ‘activated-naïve’ B cells (24), was the most abundant of all three clusters.

Together, these results demonstrate that the IgA and IgD/IgM activated compartments are heterogenous as seen for IgG. Also, in IgD/IgM B cells, absence of CD21 and expression of CD71, CD11c, or CD24 are indicators of B cell differentiation states that arise early after antigen encounter.

### The early and transient activated B cell subpopulations can also be identified by manual gating after antigen encounter

Unsupervised clustering allowed us to perform in depth characterization of the ActBC compartment and provided indications of early activated B cell populations that quickly respond to antigen encounter. For many B cell-mediated diseases, the (auto)antigen is unknown, but tracking the active B cell response is of great importance to assess disease activity, as a marker of relapse and treatment efficacy. We have previously shown by manual gating that around 20% of S-specific B cells were CD27^+^ CD11c^+^ seven days after SARS-CoV-2 vaccination (24). Therefore, we investigated if the unique phenotypic profiles of highly dynamic ActBC populations could also be employed to track active B cell responses in total CD19^+^ B cells, without the need of antigen labelling. We focused on the transient ActBC clusters 22, 25, 26 and 27 together to track the early B cell response after vaccination. Total ActBCs were manually gated from bulk CD19^+^ B cells by selecting for IgG^+^ CD27^+/lo^ CD38^lo^ CD21^-^ CD71^+^ B cells **(Figure 5A**). We were able to identify a significant expansion of IgG^+^ ActBCs at day seven after second vaccination (**Figure 5B**). To further explore this compartment, we first gated on CD11c^+^ CD24^+/-^ and CD11c^lo^ CD24^-^ to be able to detect the transient dynamic observed for clusters 25, 26 and 27 **(Figure 5A**). We found that the highly dynamic ActBC subpopulations were expanded at day seven after second vaccination and also significantly reduced six months after, indicating that this combination of markers can be used to detect activated B cell responses without the use of antigen-specific probes (**Figure 5C**). Cluster 22 also showed a transient dynamic after vaccination, so we then gated on CD27^lo^ CD45RB^+^ ActBCs and also confirmed this dynamic of early expansion and significant contraction between day seven and six months after vaccination (**Figure 5D-E**). Approximately, 38% of the IgG^+^ ActBC expansion seven days after second vaccination was S-specific compared to 0.2% at baseline and 11% six months after vaccination, indeed showing that shortly after exposure the ActBC compartment is enriched for B cells specific for the recently encountered antigen (**Figure 5F**).

**Figure 5.**
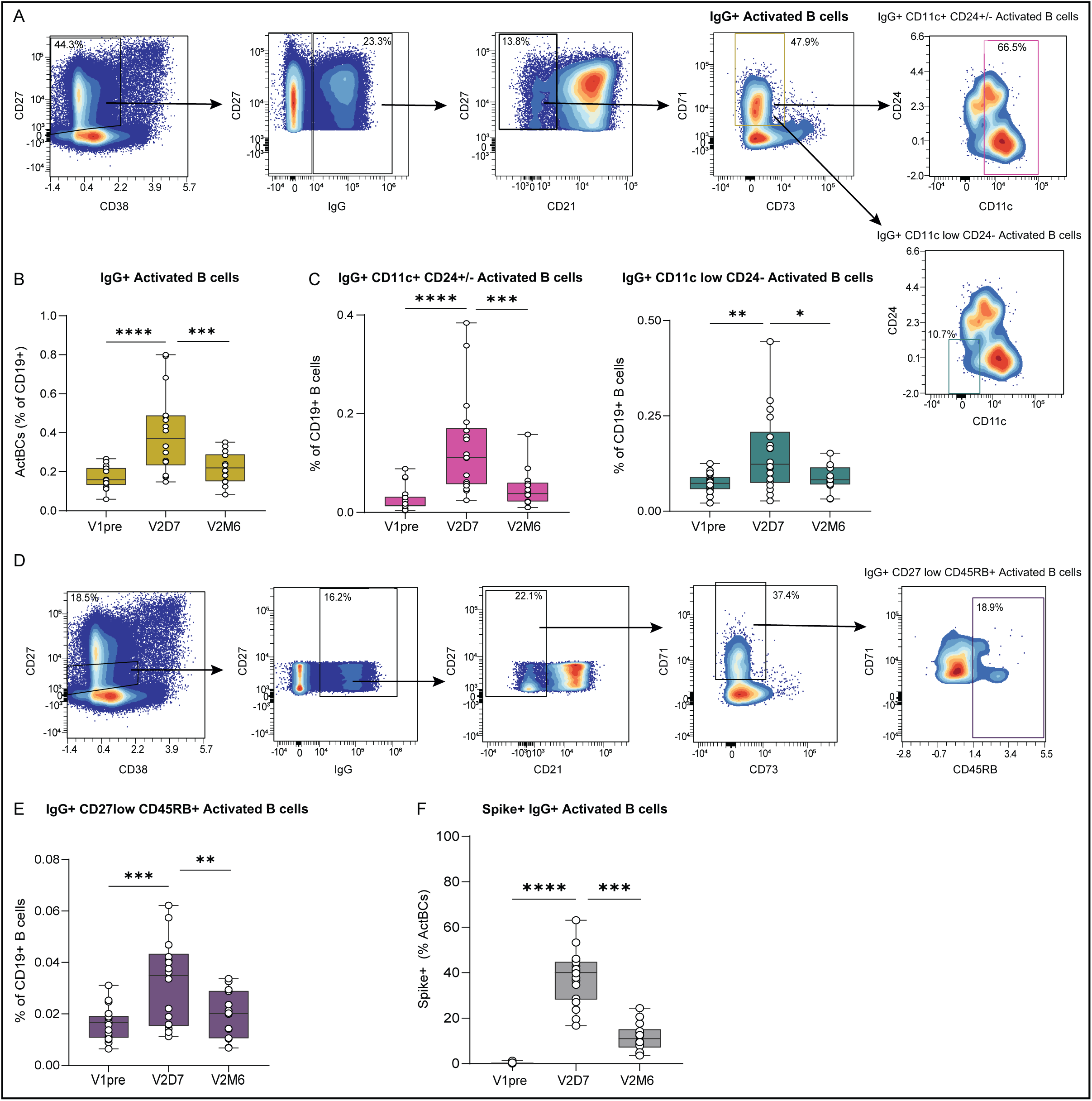
The early and transient activated B cell subpopulations can also be identified by manual gating after antigen encounter. **A.** Representative gating strategy to detect IgG^+^ ActBCs in total CD19^+^ B cells from healthy donor PBMCs at V2D7 and further CD24 and CD11c gating strategy. **B.** Proportion of ActBCs as percentage of CD19^+^ B cells. **C**. Proportion of IgG^+^ CD11c^+^ CD24^+/-^ ActBCs and IgG^+^ CD11c^lo^ CD24^-^ as percentage of CD19^+^ B cells. **D.** Representative gating strategy to detect IgG^+^ CD27^lo^ CD45RB^+^ ActBCs in total CD19^+^ B cells from healthy donor PBMCs at V2D7. **E.** Proportion of IgG^+^ CD27^lo^ CD45RB^+^ ActBCs as percentage of CD19^+^ B cells. **F**. Proportion of Spike^+^ B cells as percentage of IgG^+^ ActBCs. V1pre (n=18), V2D7 (n=18) and V2M6 (n=14). Statistical significance was assessed using Wilcoxon signed-rank test for paired data and p-values were corrected for multiple comparison using *post hoc* Bonferroni-Holm’s test. (\**P* < 0.05, \*\**P* < 0.01, \*\*\**P* < 0.001, \*\*\*\**P* < 0.0001).

Overall, our findings reveal deeper insights into the transient phenotypes of ActBC compartment. Within this compartment, we identified, in total CD19^+^ B cells, four distinct clusters that expand early and rapidly contract over time based on the marker combination of CD11c, CD24, CD45RB and CD27. These clusters represent the most promising candidates for tracking autoreactive B cells, highlighting their critical role in early immune responses.

## Discussion

Although recent advances have been made in characterizing the phenotypic diversity of the different B cell stages following antigen exposure (10, 11, 17, 24, 35, 37), better insights in the different populations that occur early after antigen exposure in the immune response will support further understanding, monitoring and possible intervention in the B cell response differentiation. In this study, we have built a comprehensive and dynamic atlas of B cell populations present at different stages during an active immune response by performing deep immune profiling of antigen-specific B cells in a longitudinal cohort study following SARS-CoV-2 vaccination in healthy individuals. We have demonstrated a higher level of complexity of the antigen-specific B cell responses than was previously acknowledged, specifically within the early arising ActBC compartment where we identified six ActBCs phenotypes, including CD11c^+^ CD71^+^ ActBCs. Furthermore, examining these populations at both an early and later timepoint after vaccination allowed us to investigate their dynamics over time, providing a deeper understanding of their contribution to an active B cell response after antigen encounter.

Classical CD27^+^ MBCs are the B cells generically used to measure the size of the memory B cell compartment after antigen-experience. They are generally defined as a homogenous population of long-lived antigen-experienced B cells. Recent advances have been made, however, driven by high-throughput approaches, in defining different subphenotypes of circulatory MBCs (9–11). Characterizing the different MBC subphenotypes and their longevity holds great value, especially for optimizing vaccination responses and for detecting undesired MBC responses as in the context of autoimmunity. Both S-specific IgG^+^ and IgA^+^ CD21^+^ CD73^+^ CD45RB^+^ MBCs were present most prominently six months following vaccination. In addition, it was the most dominant IgG^+^ B cell cluster observed for HA-, TT- and RSV-F-specific antigens that were encountered longer ago in our cohort at this time of sampling. This suggests that the CD21^+^ CD73^+^ CD45RB^+^ marker combination for CD27^+^/IgG^+^ MBCs characterize a truly long-lived resting MBC compartment (11,35). MBCs expressing CD45RB, CD73 and CD95 have recently been defined as the most ‘advanced’ naïve-distal MBC population (35). Functionally, CD73 is used as a marker for metabolic quiescence in naïve B cells and might be relevant for long-lived properties of resting MBCs in circulation as well (37). Our data also showed CD27^lo^ CD21^+^ CD73^-^ CD45RB^-^ MBCs to be strongly expanded six months after vaccination, however they did not contribute to the long-lasting memory compartment of previously encountered antigens. This cluster resembles CD27^-^ CD21^+^ resting MBCs observed by others one year after Influenza and SARS-CoV-2 vaccination (10,38), and may reflect a more immature, less long-lived MBC population. Lastly, the smallest observed MBC population (CD21^-^ CD73^-^ CD45RB^+^, cluster 14) might resemble the naïve-proximal MBC population also described by McGrath and colleagues given its expression of CD45RB and absence of CD73 (9). However, the low expression of CD21, together with a less profound expansion between day seven and six months after vaccination, suggests a more complex phenotype and may indicate that this cluster is an intermediate population between ActBCs and MBCs. Together, our data and that of others show that the CD21^+^ CD73^+^ CD45RB^+^ marker combination, seems to be the phenotype that marks long-lived IgG^+^ MBCs.

Upon identification of the early B cell compartment, we found that IgG^+^ ActBCs dominated the S-specific response, together with ASCs and DN2 (CD27^-^ CD21^-^ CD11c^+^) at day seven after vaccination, consistent with findings reported by others (14, 20, 37, 40). ActBCs have been identified as GC-experienced B cells, which are present early after antigen exposure (11–14). Inconsistencies between different studies in defining and naming ActBCs has made comparisons between populations challenging, ActBCs are generally defined as CD27^+^ CD21^-^ B cells (11–14), while others refer to them as activated MBCs (15, 37, 39, 41). Here, we defined the ActBC compartment as CD27^+/lo^ CD38^lo^ CD21^lo^ CD71^+^ CD95^hi^ B cells, combining absence of CD21 with expression of CD71 and CD95 as also employed by others (11, 12, 14). From this definition and from our clustering analysis, CD71 was the best discriminative marker to capture the ActBC compartment. ASCs also express CD71, but this compartment can be discriminated by being CD19^-/lo^ and no longer in B cell stage. Notably, we found that the ActBC compartment was CD38^lo^ while others define it as CD38^+^ (14, 37). This likely can be explained by the fact that these studies looked at the relative CD38 expression within the B cells, excluding the CD38^hi^ ASCs. In our study, all CD38^hi^ CD71^+^ were confined to the ASC compartment.

Deep profiling of the IgG^+^ ActBC compartment allowed us to subdivide it into six clusters based on differential expression of certain markers and distinct dynamics, together providing a higher resolution of the ActBC compartment. The six clusters could dynamically be divided in two groups, with both groups arising early after antigen exposure, but one group contracting faster than the other in the long term. Clusters 23 and 24 arose early after antigen encounter, but did not contract in all individuals over time. It is tempting to speculate that the prolonged presence of these clusters six months after vaccination may suggest that they were induced during a longer period in GC-reactions, in line with the persistent GC after mRNA vaccination and the suggested GC-experienced nature of IgG^+^ ActBCs in general (12, 13 ,34, 42).

The early arising ActBC clusters 22, 25, 26 and 27 contracted between day seven and month six after vaccination One of our most striking findings was co-expression of CD71 and CD11c in the IgG^+^ ActBC clusters 25 and 27. Indeed, our previous findings revealed by manual gating that around 20% of S-specific B cells were CD11c-expressing CD27^+^ CD38^-^ B cells, referred to as age-associated (atypical) B cells, seven days after SARS-CoV-2 vaccination, while CD71 expression was not regarded (24). IgG^+^ DN2, which lack CD27 and CD21 expression but express CD11c, also arose early and contracted to pre-vaccination levels six months after vaccination. DN2 are thought to be part of the EF B cell response and have been extensively studied in the context of chronic infections and autoimmunity (19, 43). Current unsupervised analysis and previous manual gating (24) indicate that besides ActBCs, DN2 cells are also a relevant antigen-specific compartment of the active vaccination response. The similar dynamics of the CD11c^+^ ActBCs with DN2 and the finding that they contract more in comparison to the other ActBC (clusters 23, 24) populations, may indicate that CD11c^+^ ActBCs and DN2 could share an EF origin. However, the similar dynamics in contraction with the CD11c^-^ ActBC cluster 22 does not support that hypothesis. This unclarity is underscored by a recent study showing plasticity between human CD21^+^ CD27^+^ resting and CD21^-^ CD27^+^ activated MBCs and DN2 (CD21^-^ CD27^-^) during SARS-CoV-2 recall response, also revealing clonality relationships between the different MBC subsets and DN2 (15). Future analysis of the level of SHM and clonal relationship will shed light in the relatedness of these populations. All in all, the position of CD11c^+^ CD71^+^ ActBC clusters in EF- or GC-derived B cell differentiation remains to be elucidated by further research. Future analysis of the level of SHM and clonal relationships between these clusters and the long-lived MBCs are needed to define how these populations relate to each other and which clusters are precursor stages of GC-derived long-lived MBC.

In addition to investigating the relationships between ActBCs, long-lived MBCs and EF B cells, several studies have aimed to address whether certain subsets of ActBCs contribute to the resting MBC pool or are poised for transition to ASC lineage (12, 13, 37). In our data, in the ASC compartment, cluster 36 might represent a pre-ASC B cell stage, as it still expressed CD20 and was located in a phenotypic continuum between IgG^+^ ActBCs and IgG^+^ ASCs in the UMAP. In addition, this cluster was high in CD30 expression, in line with recent assignments of CD30 expression to a pre-plasmablast GC-derived populations (16, 36, 44). It is important to highlight that cluster 36 is in a phenotypic continuum with the DN3 cluster and the IgG^+^ ActBC cluster 24. As these clusters have lower expression of CD20 compared to the other ActBCs, they may represent good candidates predisposed for ASC differentiation. Further research on isolated subsets involving scRNAseq and analyses of clonal evolution over time are needed to identify the decision point between MBCs and ASCs.

Finally, our data demonstrate that an increase in transient IgG^+^ ActBC clusters early after antigen exposure can be detected in the total CD19^+^ B cell compartment by gating on IgG^+^ CD27^+/lo^ CD38^lo^ CD21^-^ CD71^+^ B cells in combination with CD11c, CD24 and CD45RB. Detection of early activated B cells, correlating with recent antigen exposure, without the use of antigen-specific probes offers the potential to monitor recently activated B cells with unknown antigen specificity. For many detrimental B cell responses, for instance in the context of autoimmunity, the autoantigens that drive the B cell response are unknown. This hampers investigation and identification of the autoreactive B cell compartments. Consequently, monitoring of the autoreactive B cell compartment before therapy and effects of therapy on this compartment cannot be executed. As the right gating strategy now allows the identification of transient IgG^+^ ActBC clusters in total B cells, it now becomes feasible to assess if these clusters can be used as a proxy for autoreactive B cells having recently encountered auto-antigen, for instance in the case of autoimmune diseases of acute onset like myasthenia gravis or encephalomyelitis. Next steps would be to assess if effective therapy would diminish the size of this compartment and if so, if the specific transient IgG^+^ ActBC clusters would also have potential to identify patients prone to relapses before onset of actual clinical symptoms.

To conclude, this study provides a detailed and dynamic map of the antigen-specific B cell compartment after antigen exposure. The revelation of different ActBCs clusters with different phenotypes and contraction dynamics, brings us closer to understanding B cell differentiation trajectories towards either MBCs or ASCs. The possibility of tracking an active immune response without the use of antigen labeling offers new opportunities to explore early B cell differentiation in response to unknown antigens as in the context of autoimmune diseases. This information lays the framework for future research on identifying factors regulating human B cell differentiation. Further investigation of the ActBC compartment may allow identification of long-lived ASC or MBC precursors that could be potentially targeted in the context of unwanted antibody formation or for improving vaccination strategies against existing and emerging pathogens.

## Author contributions

AtB and SMvH led the study. SMvH is the guarantor. SMvH and TWK supervised the study. FE, TWK and SMvH conceptualized the study. SMvH, AtB, LFB, LHK, AB, NJMV, MC, MJvG, JJGV, TR and MS designed the methodology. LHK, LYLK, AB, LFB, NJMV, MCD, TJ and GK performed the experiments. LFB, LHK, LYLK, SMvH and AtB verified the overall replication/reproducibility of the research output. LFB, LHK, LYLK, AB and NJMV analyzed the data. LYLK, KPJvD, EWS, LB, GJW and SWT provided study materials. LYLK, KPJvD, EWS and LW managed the research data. LYLK, KPJvD, EWS, LW, TR, TWK, FE, SMvH and AtB managed and coordinated the research activity and planning. TWK, FE and SMvH acquired the financial support for the project leading to this publication. LYLK, LHK, MC and NJMV designed and implemented computer codes. LFB visualized the work. LFB, SMvH and AtB wrote the original draft. All authors reviewed and approved the manuscript.

## Acknowledgements

We would like to thank all donors who participated in the study, the Sanquin COVID-19 cryo and biobank facility for processing of samples. Also, we would like to thank Cora Chadick and Liesbeth Paul from the VUmc Core facility for providing technical assistance with the spectral flow cytometry experiments. Further, we will like to thank Jim Keijser, Sophie Keijzer and Olvi Cristianawati from Sanquin for performing the ELISAs. We would like to acknowledge the funding provided by ZonMw for this study and thank all our T2B partners, including the patient groups, as well as Health Holland for their contributions and support during this research.

## Funding

This research project was supported by ZonMw (The Netherlands Organization for Health Research and Development, #10430072010007). LHK and MCD received funding for this study from Sanquin Blood Supply program grant PPOC OPTIMAL, project number L2506.

## Ethics approval

This study was approved by the medical ethical committee (NL74974.018.20 and EudraCT 2021-001102-30, local METC number: 2020_194). Participants gave informed consent to participate in the study before taking part.

## Competing interests statement

FE, GW, SMvH and TWK report (governmental) grants from ZonMw to study immune response after SARS-CoV-2 vaccination in auto-immune diseases. FE also reports grants from Prinses Beatrix Spierfonds, CSL Behring, Kedrion, Terumo BCT, Grifols, Takeda Pharmaceutical Company, and GBS-CIDP Foundation; consulting fees from UCB Pharma and CSl Behring; honoraria from Grifols. All other authors report no competing interests to this manuscript.

## SUPPLEMENTARY MATERIAL

**Supplementary figure 1. A.** Anti-RBD IgG levels for all 18 individuals at V1pre, V1D28, V2pre, V2D7, V2D28, V2M6. **B.** Gating strategy for the detection of CD19^+^ B cells. **C.** Gating strategy for the detection of S-specific B cells using a combinatorial probe staining. S-specific B cells are detected as double positive for the binding of the same antigen combined with two different fluorophores. RBD-specific B cells are detected from S-specific B cells. **D.** Percentage of S/RBD-ratio of total CD19^+^ B cells at V2D7 (n=18) and V2M6 (n=14) **E.** UMAP representation showing normalized expression of all markers in all antigen-specific B cells. **F.** UMAP projection of RBD-specific B cells per time point. **G.** Stacked bar graph representing the mean proportion of RBD^+^ B cells (% of CD19^+^) for the 16 main populations per time point. Statistical significance was assessed using Wilcoxon signed-rank test for paired data and p-values were corrected for multiple comparison using *post hoc* Bonferroni-Holm’s test. (\**P* < 0.05, \*\**P* < 0.01, \*\*\**P* < 0.001, \*\*\*\**P* < 0.0001).

**Supplementary figure 2. Expression pattern of IgG^+^ ActBC, DN B cells and MBC clusters.** Analysis of cell surface expression by histogram representation of 14 relevant markers in IgG ActBC, DN B cell and MBC clusters.

