## Supplementary figures and images for "Characterization of a heterogenous activated B cell compartment arising early after antigen exposure preceding long-lived memory B cell formation"

### Supplementary figure 1

Supplementary Figure 1

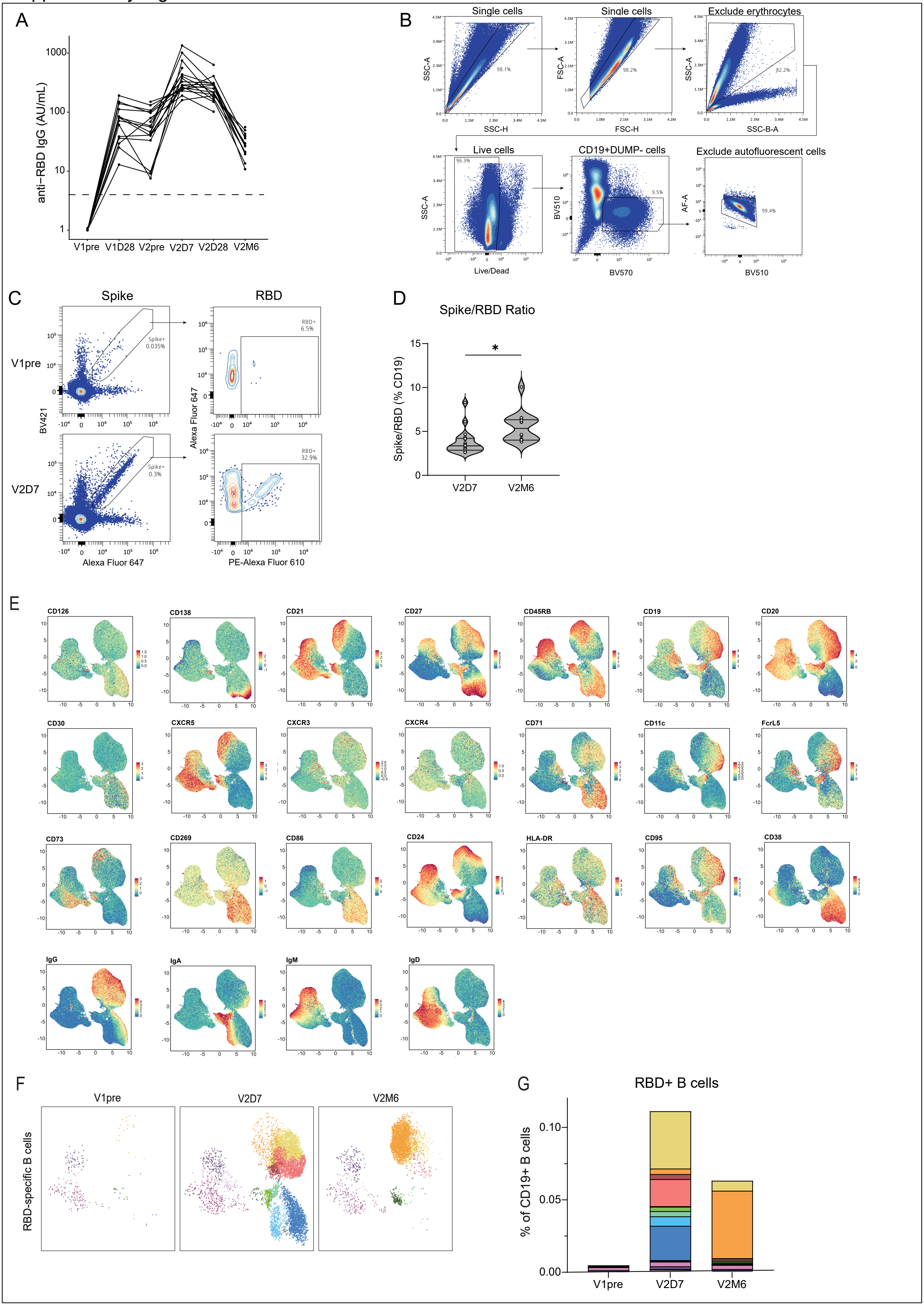

### Supplementary figure 2

Supplementary Figure 2

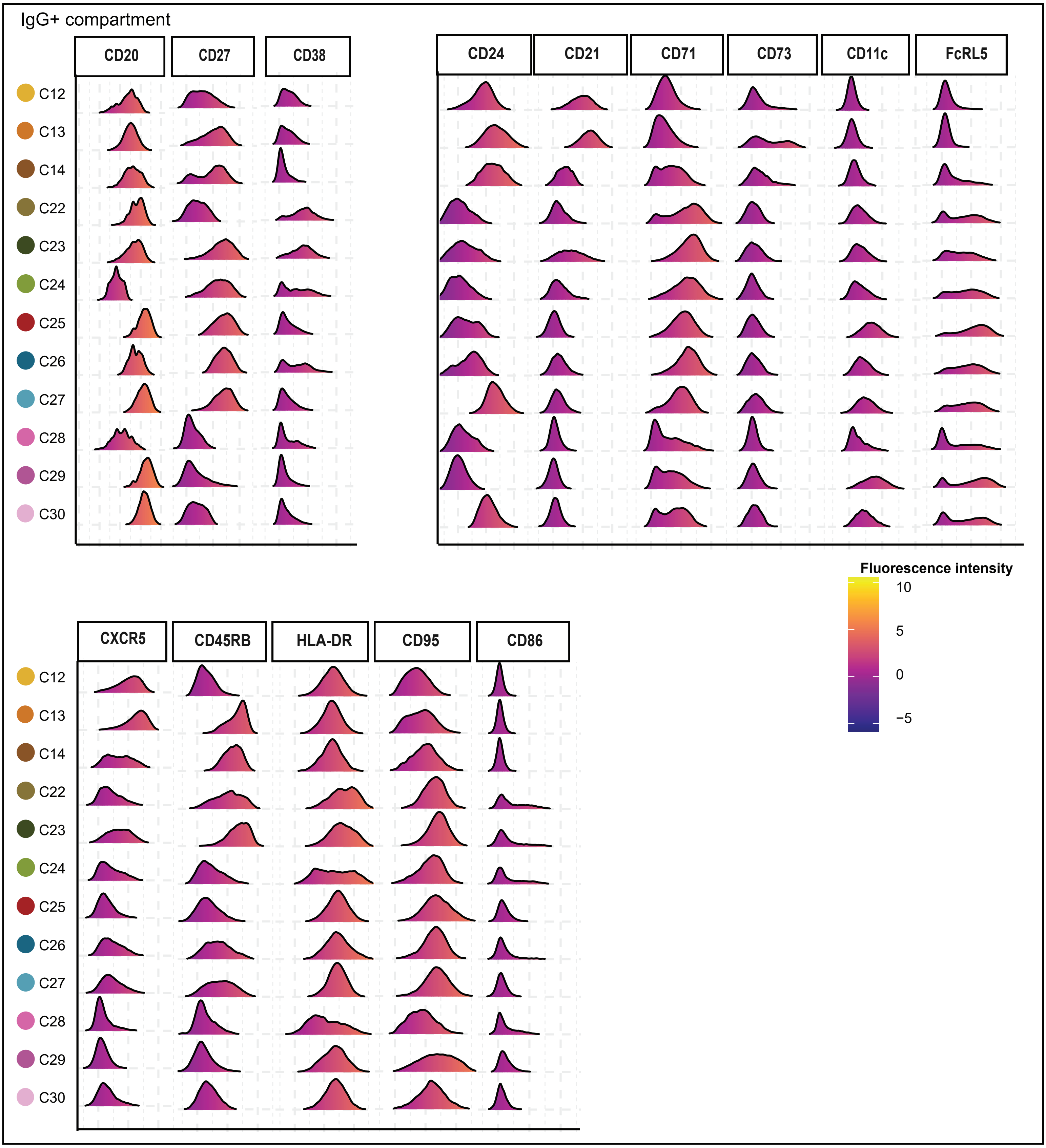
