## Supplementary tables 1_2_3 for "Characterization of a heterogenous activated B cell compartment arising early after antigen exposure preceding long-lived memory B cell formation"

**Supplementary table 1: Panel design antigen probes**

| **Protein antigen** | **Fluorochrome-conjugated streptavidin** | **Concentration** | **Source** | **Catalog No** |
| --- | --- | --- | --- | --- |
| Wuhan  SARS-CoV-2 Spike | Alexa Fluor® 647 | 0.5 ug/ul | BioLegend | 405237 |
|  | Brilliant Violet™ 421 | 0.5 ug/ul |  | 405225 |
| Wuhan  SARS-CoV-2 RBD | PE-Alexa Fluor® 610 | 1 ug/ul | Invitrogen | S20982 |
| H1N1 HA | BUV615 | 0.1 ug/ul | BD | 613013 |
|  | Brilliant Violet™ 421 | 0.5 ug/ul | BioLegend | 405225 |
| Nucleocapsid | Brilliant Blue™ 515 | 0.1 ug/ul | BD | 564453 |
|  | Alexa Fluor® 647 | 0.5 ug/ul | BioLegend | 405237 |
| RSV F | BUV615 | 0.1 mg/mL | BD | 613013 |
|  | Alexa Fluor® 647 | 0.5 ug/ul | BioLegend | 405237 |
| Tetanus toxoid | Brilliant Blue™ 515 | 0.1 ug/ul | BD | 564453 |
|  | Brilliant Violet™ 421 | 0.5 ug/ul | BioLegend | 405225 |

**Supplementary table 5: B cell cluster annotations**

Laura F will make this

**Supplementary table 2: B cell panel**

| **Antibody** | **Clone** | **Fluorochrome** | **Dilution*** | **Source** | **Catalog no** |
| --- | --- | --- | --- | --- | --- |
| CD24 | ML5 | BUV395 | 1/80 | BD | 563818 |
| Viability | - | Live/Dead Blue | 1/1000 | Thermo Fisher | L23105 |
| HLA-DR | G46-6 | BUV496 | 1/80 | BD | 749866 |
| IgD | IA6-2 | BUV563 | 1/160 | BD | 741394 |
| CD126 | M5 | BUV661 | 1/20 | BD | 752527 |
| CD138 | MI15 | BUV737 | 1/100 | BD | 612834 |
| CD21 | B-ly4 | BUV805 | 1/160 | BD | 742008 |
| CD20 | 2H7 | cFluor V450 | 1/40 | Cytek | SKU R7-20015 |
| CD27 | L128 | BV480 | 1/20 | BD | 566188 |
| CD3 (DUMP) | UCHT1 | BV510 | 1/50 | BioLegend | 300448 |
| CD4 (DUMP) | OKT4 | BV510 | 1/50 | BioLegend | 317444 |
| CD16 (DUMP) | 3G8 | BV510 | 1/50 | BioLegend | 302048 |
| CD56 (DUMP) | HCD56 | BV510 | 1/50 | BioLegend | 318340 |
| CD19 | HIB19 | BV570 | 1/40 | BioLegend | 302236 |
| CD11c | B-ly6 | BV605 | 1/160 | BD | 563929 |
| CXCR5 | RF8B2 | BV650 | 1/20 | BD | 740528 |
| IgM | MHM-88 | BV711 | 1/80 | BioLegend | 314540 |
| CD71 | M-A712 | BV750 | 1/80 | BD | 747308 |
| FcRL5 | 509F6 | BV785 | 1/100 | BD | 749602 |
| CD73 | AD2 | cFluor B532 | 1/80 | Cytek | SKU R7-20017 |
| CXCR3 | 1C6 | BB700 | 1/25 | BD | 566532 |
| CXCR4 | 12G5 | PerCP-eFluor710 | 1/50 | Thermo Fisher | 46-9999-42 |
| CD45RB | MEM-55 | PE | 1/80 | BioLegend | 310204 |
| IgG | G18-145 | PE-CF594 | 1/160 | BD | 562538 |
| CD95 | DX2 | PE-Cy5 | 1/320 | BioLegend | 305610 |
| CD30 | BY88 | PE-Cy7 | 1/12 | BioLegend | 333918 |
| CD269 | 19F2 | APC | 1/25 | BioLegend | 357506 |
| CD86 | 2331 (FUN-1) | APC-R700 | 1/320 | BD | 565149 |
| IgA | IS11-8E10 | APC-Vio770 | 1/320 | Miltenyi Biotec | 130-113-473 |
| CD38 | HIT2 | APC-Fire810 | 1/160 | BioLegend | 303550 |

*Dilutions used were optimized in-house, these should be used as a guideline and optimized for individual laboratories.

**Supplementary table 3: B cell cluster annotations**

| Cluster | B cell subpopulation | B cell population | Isotype | Defining markers | Additional markers | Other markers |
| --- | --- | --- | --- | --- | --- | --- |
| 1 | IgD low CD73+ HLA-DR+ naive B cell | Naive B cell | IgD | CD19+ CD20+ CD27- CD38- | CD21+ CD73++ HLA-DR++ | CXCR5+ |
| 2 | IgD/ IgM transitional B cell | Transitional B cell | IgD/ IgM | CD19+ CD20+ CD27- CD38lo | CD21+ CD24+ | CXCR5+ |
| 3 | IgD/IgM CD73+ naïve B cell | Naïve B cell | IgD/ IgM | CD19+ CD20+ CD27- CD38- | CD21+ CD24+ CD73+ | CXCR5+ CXCR4+ |
| 4 | IgD CD73+ naïve B cell | Naïve B cell | IgD | CD19+ CD20+ CD27- CD38- | CD21+ CD24+ CD73+ | CXCR5+ CXCR4+ |
| 5 | IgM CD45RB+ MBC | IgM MBC | IgM | CD19+ CD20+ CD27+ CD38- | CD21+ CD24+ CD45RB+ | CXCR5+ |
| 6 | IgM CD27low CD45RB+ MBC | IgM MBC | IgM | CD19+ CD20+ CD27lo CD38- | CD21+ CD24+ CD45RB+ | CXCR5+ |
| 7 | IgD CD73low CD45RB+ naïve B cell | CD45RB+ naïve B cell | IgD | CD19+ CD20+ CD27- CD38- | CD21+ CD24+ CD73lo CD45RB+ | CXCR5+ |
| 8 | IgD/IgM CD73low CD45RB+ naïve B cell | CD45RB+ naïve B cell | IgD/ IgM | CD19+ CD20+ CD27- CD38- | CD21+ CD24+ CD73lo CD45RB+ | CXCR5+ |
| 9 | IgA CD45RB- CD73- MBC | IgA MBC | IgA | CD19+ CD20+ CD27+ CD38- | CD21+ CD24lo CD73- CD45RB- | CXCR5+ CD11c- |
| 10 | IgA CD45RB+ CD73+ MBC | IgA MBC | IgA | CD19+ CD20+ CD27+ CD38- | CD21+ CD24+ CD73+ CD45RB+ | CXCR5+ CD11c- |
| 11 | Undefined | Not BC | - | - | - | - |
| 12 | IgG CD21+ CD73- MBC | IgG MBC | IgG | CD19+ CD20+ CD27lo CD38- | CD21+ CD24+ CD73- CD45RB- | CXCR5+ |
| 13 | IgG CD21+ CD73+ CD45RB+ MBC | IgG MBC | IgG | CD19+ CD20+ CD27+ CD38- | CD21+ CD24+ CD73+ CD45RB+ | CXCR5+ |
| 14 | IgG CD21low CD73- CD45RB+ MBC | IgG MBC | IgG | CD19+ CD20+ CD27+ CD38- | CD21lo CD24+ CD73- CD45RB+ | CXCR5- |
| 15 | IgA CD45RB+ CD24- Activated B cell | IgA Activated B cell | IgA | CD19+ CD20+ CD27+ CD38- CD71hi | CD24- CD45RB+ | CXCR3+  CXCR5- |
| 16 | IgA CD45RB- CD24+ CD11c+ Activated B cell | IgA Activated B cell | IgA | CD19+ CD20+ CD27+ CD38- CD71hi | CD24+ CD45RB- CD11c+ | CXCR3+ CXCR5- |
| 17 | IgA DN2 | IgA DN2 | IgA | CD19+ CD20+ CD27- CD38- CD11c hi | CD71- CD24- CD95+ | CXCR3+ CXCR5- |
| 18 | IgD/IgM CD24+ MBC | IgD/IgM MBC | IgD/ IgM | CD27+ CD21- | CD71- CD24+ CD95+ CD45RB- | CXCR3+ CXCR5- |
| 19 | IgD/IgM CD21- CD24+ Activated B cell | IgD/IgM Activated B cell | IgD/ IgM | CD19+ CD20+ CD27- CD38- | CD21- CD71- CD24+ | CXCR3+ |
| 20 | IgM CD21- CD71+ Activated B cell | IgD/IgM Activated B cell | IgM | CD19+ CD20+ CD27- CD38- | CD21- CD71+ CD24- | CXCR3+ |
| 21 | IgD CD21- CD11c+ Activated B cell | IgD/IgM Activated B cell | IgD | CD19+ CD20+ CD27- CD38- | CD21- CD11c+ CD71- CD24- |  |
| 22 | IgG CD27low CD45RB+ Activated B cell | IgG Activated B cell | IgG | CD19+ CD20+ CD27lo CD38- CD71hi | CD24- CD95+ CD45RB+ HLA-DR+ |  |
| 23 | IgG CD45RB+ Activated B cell | IgG Activated B cell | IgG | CD19+ CD20+ CD27+ CD38- CD71hi - | CD24- CD95+ CD45RB+ HLA-DR+ |  |
| 24 | IgG CD11c low CD24- CD20low Activated B cell | IgG Activated B cell | IgG | CD19+ CD20+ CD27+ CD38- CD71hi | CD24- CD95+ CD45RB- HLA-DR lo | CD11c lo |
| 25 | IgG CD11c+ CD24- Activated B cell | IgG Activated B cell | IgG | CD19+ CD20+ CD27+ CD38- CD71hi | CD11c+ CD24- CD95+ CD45RB- HLA-DR+ |  |
| 26 | IgG CD11c low CD24- Activated B cell | IgG Activated B cell | IgG | CD19+ CD20+ CD27+ CD38- CD71hi | CD24- CD95+ CD45RB- HLA-DR+ | CD11c lo |
| 27 | IgG CD11c+ CD24+ Activated B cell | IgG Activated B cell | IgG | CD19+ CD20+ CD27+ CD38- CD71hi | CD11c+ CD24+ CD95+ CD45RB- HLA-DR+ |  |
| 28 | IgG DN3 | IgG DN3 | IgG | CD27- CD21- CD20lo | CD24- CD71- CD95+ |  |
| 29 | IgG DN2 | IgG DN2 | IgG | CD19+ CD20+ CD27- CD38- CD11c hi | CD24- CD95+ |  |
| 30 | IgG CD24+ DN2 | IgG DN2 | IgG | CD19+ CD20+ CD27- CD38- CD11c hi | CD24+ CD95+ |  |
| 31 | IgA Plasma cell | IgA ASC | IgA | CD19lo CD20lo CD27hi CD38hi CD138+ | CD86+ CD269+ CD45RB+ |  |
| 32 | IgA Plasmablast | IgA ASC | IgA | CD19lo CD20lo CD27hi CD38hi | CD86+ CD269+ CD45RB+ |  |
| 33 | IgM Plasmablast | IgM ASC | IgM | CD19lo CD20lo CD27hi CD38hi | CD86+ CD269+ CD45RB+ |  |
| 34 | IgG Plasmablast | IgG ASC | IgG | CD19lo CD20lo CD27hi CD38hi | CD86+ CD269+ CD45RB+ |  |
| 35 | IgG Plasma cell | IgG ASC | IgG | CD19lo CD20lo CD27hi CD38hi CD138+ | CD86+ CD269+ CD45RB+ |  |
| 36 | IgG CD20+ CXCR4+ Early Plasmablast | IgG ASC | IgG | CD19lo CD20+ CD27lo CD38hi | CD86+ CD269+ CD30+ | CXCR4+ |
| 37 | IgG Early Plasmablast | IgG ASC | IgG | CD19lo CD20lo CD27lo CD38hi | CD86+ CD269+ |  |
| 38 | IgA Early Plasmablast | IgA ASC | IgA | CD19lo CD20lo CD27lo CD38hi | CD86+ CD269+ |  |

**Abbreviations:** BC, B cell; DN, double negative; MBC, memory B cell; PB, plasmablast; PC, plasma cell.
